# EGO-like complex regulates TOR (Target of Rapamycin) activity and localization in *Neurospora*

**DOI:** 10.64898/2025.12.05.692677

**Authors:** Rosa Eskandari, Maryam Fayyazi, Patricia Lakin-Thomas

## Abstract

The TOR (Target of Rapamycin) signalling pathway is found in all eukaryotes and integrates nutrient and stress signals to control cell growth. It is well-studied in yeast and mammals but is less well understood in filamentous fungi. We previously identified TOR pathway components VTA (homologous to the vacuole-bound EGO complex of yeast) and GTR2 (homologous to Rag GTPases) as essential to maintaining circadian rhythms in the filamentous fungus *Neurospora crassa*. Therefore we are interested in defining the TOR pathway and its regulation in *N. crassa*. In yeast and mammals, TOR kinase is activated by carbon sources and amino acids. We report here that on high glucose medium, TOR is insensitive to added amino acids. On low glucose, TOR is activated by added glucose and amino acids. VTA and GTR2 knockouts block the activation of TOR by amino acids but not by glucose, identifying their function in an amino acid-sensing pathway. Live cell microscopy of KOG1 (a component of TOR complex 1) and GTR2 localizes them to punctate bodies near the vacuole. This localization is lost and the proteins are largely cytoplasmic under starvation conditions, in the presence of TOR inhibitor Torin II, and in the VTA knockout. This indicates that VTA acts as the vacuolar anchor for activated TOR complex. Co-immunoprecipitation of KOG1-FLAG and GTR2-FLAG confirms their cytoplasmic localization in VTA knockout and identifies TORC1 complex components TCO89 and LST8. These results focus attention on amino acid sensing through VTA and GTR2 as potentially regulating circadian rhythmicity in *N. crassa*.

## 1. Introduction

TOR (Target of Rapamycin) is a nutrient sensitive central controller of cell growth in eukaryotes and is the target of the drug rapamycin. The mammalian target of rapamycin (mTOR) integrates both extracellular and intracellular signals to promote cell growth, proliferation, and anabolic metabolism (Saxton and Sabatini, 2017). TOR has also been well-studied in yeast (Loewith and Hall, 2011) but much less is known about TOR in filamentous fungi. Our laboratory studies the filamentous fungus *Neurospora crassa*, and we are interested in describing the function and regulation of TOR in our model organism.

TOR is a Ser/Thr protein kinase that is found in two complexes, TORC1 and TORC2 (Laplante and Sabatini, 2012). TORC1 is the rapamycin-sensitive complex that controls growth and metabolism in all eukaryotes (Liu and Sabatini, 2020). TORC2 is rapamycin insensitive, is less well-understood and has a more evolutionarily divergent set of functions (Thorner, 2022). The presence of growth hormones (in mammals) and of nutrients causes activation of TORC1, which leads to an increase in anabolic processes, such as protein synthesis and mRNA synthesis through phosphorylation of downstream substrates. In contrast, starvation or TORC1 inhibitors such as rapamycin lead to decreased TORC1 activity and a drop in substrate phosphorylation. Inhibition of TORC1 causes cells to switch from anabolic to catabolic metabolism and enter a quiescent state (Hughes Hallett et al., 2014).

In mammals and yeast, the mTORC1/TORC1 complex includes three essential components: Raptor/ Kog1, mLST8/Lst8, and mTOR/Tor; yeast TORC1 also includes the fungal-specific subunit Tco89 (Tafur et al., 2020). The presence of Kog1/Raptor is necessary to recruit substrates to the TOR kinase and it acts a regulator for TOR activity. mLst8/Lst8 is assumed to stabilize the complex (Saxton and Sabatini, 2017). Amino acid and nitrogen signals act through a complex that contains small GTPases known as Gtr1/2 in yeast and RagA/C in mammals. These complexes are localized on the vacuolar or lysosomal membrane, and mediate and respond to these signals by binding and activating TORC1 (Sancak et al., 2010). At the lysosomal/vacuolar membrane, the GTPases are anchored to the membrane by the Ragulator complex (in mammals) or the EGO complex (in yeast). During energy starvation TOR is inhibited through the AMPK pathway. Glucose deprivation leads to AMPK activation and the phosphorylation of the Kog1/Raptor component of TORC1 in yeast and mammals, thereby inhibiting TORC1 activity (Smiles et al., 2024).

Our lab’s investigation of TOR derives from our central interest in the molecular mechanism of the circadian clock in *N. crassa.* Circadian (daily) rhythmicity is a fundamental property of cellular physiology in most eukaryotes as well as some bacteria, and *N. crassa* has served as a unique model organism for many years for investigating circadian rhythmicity. The production of asexual conidiospores (conidiation) is rhythmically controlled by the circadian clock and serves as a convenient readout of the state of the circadian oscillator. We have previously reported (Eskandari et al., 2021; Li et al., 2011; Ratnayake et al., 2018) that two components of the TOR pathway, VTA (homologous to a yeast EGO complex protein) and a binding partner of VTA, GTR2 (homologous to yeast Gtr2) are required to maintain normal circadian rhythmicity in *N. crassa*. We have recently reported (Akhtari et al., 2025) that TORC1 activity is rhythmic in *N. crassa* and is closely coupled to the circadian conidiation rhythm. Our goal is to determine what makes TORC1 activity rhythmic, how the *vta^ko^* and *gtr2^ko^* knockout mutations affect that rhythmicity, and what role the TOR pathway plays in the circadian clock system. To approach that goal, we first need to describe the regulation of TOR activity in *N crassa*.

In the present study, we have assayed the effects of *vta^ko^*and *gtr2^ko^* on the regulation of TOR activity by carbon source (glucose) and amino acids. To investigate the localization of the TORC1 complex, we have used strains carrying fluorescently tagged GTR2 and KOG1 proteins in the presence and absence of activating nutrients, the TOR inhibitor Torin II, and in the *vta^ko^*background. The composition of TOR pathway complexes was investigated by co-immunoprecipitation of FLAG-tagged KOG1 and GTR2. Our findings point to a role for VTA and GTR2 in amino acid sensing but not carbon source response, and implicate vacuolar localization of TORC1 as essential for amino acid regulation of TOR activity and maintenance of circadian rhythmicity.

## 2. Materials and methods

### 2.1 Strain construction

All strains utilized in this paper carry the *ras^bd^*, *csp-1* and *chol-1* mutations. “Wild type” indicates a *csp-1; chol-1 ras^bd^* strain with no additional mutations, with or without S6-FLAG as indicated. The *ras^bd^*mutation (Belden et al., 2007) makes conidiation resistant to the repressing effect of carbon dioxide, allowing the development of rhythmic conidiation bands in closed cultures such as race tubes. The CSP-1 gene product is a transcriptional repressor (Sancar et al., 2015); the *csp-1* mutation is used in our lab for convenience to prevent conidial separation, reducing self-contamination of cultures. Strains carrying the *chol-1* mutation require choline for normal growth and rhythmicity (Lakin-Thomas, 1996); all culture media in this paper contain a sufficient concentration of choline to repair the defect in *chol-1* strains.

The *vta^ko^*, *gtr2^ko^*, VTA-GFP and GTR2-FLAG strains were constructed as previously reported (Eskandari et al., 2021; Ratnayake et al., 2018). The *kog1* gene in *N. crassa* was previously annotated in the FungiDB database (https://fungidb.org/fungidb/app) and was identified as NCU00621, TORC1 growth control complex subunit Kog1. The KOG1-GFP, KOG1-FLAG and GTR2-RFP plasmids were constructed using recombination-mediated plasmid construction in *Saccharomyces cerevisiae* to make knock-in cassettes (Honda and Selker, 2009), as previously described (Eskandari et al., 2021). Primers used to amplify the coding sequences from *N. crassa* genomic DNA and construct the knock-in plasmids are listed in Table 1. KOG1-GFP, KOG1-FLAG, and GTR2-RFP plasmids were extracted from yeast and amplified in *E. coli*. Knock-in cassettes were amplified from bacterial plasmids using 5’ forward and 3’ reverse specific primers, and the PCR products were then transformed by electroporation into the *mus-51 his-3* strain of *N. crassa* deficient in non-homologous end joining (Honda and Selker, 2009). Transformed colonies were picked and grown on slants. To confirm the presence of the cassette, spores were genotyped using the Terra Direct PCR kit (Takara Bio), and *gtr2* and *kog-1-*specific primers. Heterokaryon strains were crossed into our lab strain (*csp-1; chol-1 ras^bd^*; Δ*vta)*. Around 800-1000 progeny were investigated from each cross. Slant cultures were genotyped for the presence and absence of *csp-1, his, chol-1, ras^bd^*, and *mus-51.* gDNA extraction was performed on candidates, and the presence of desired cassettes and the *vta* mutation were investigated by PCR. Homokaryons were obtained from GTR2-RFP, KOG1-GFP and KOG1-FLAG strains. KOG1-FLAG protein expression was confirmed by Western blotting using anti-FLAG antibody.

**Table 1.**
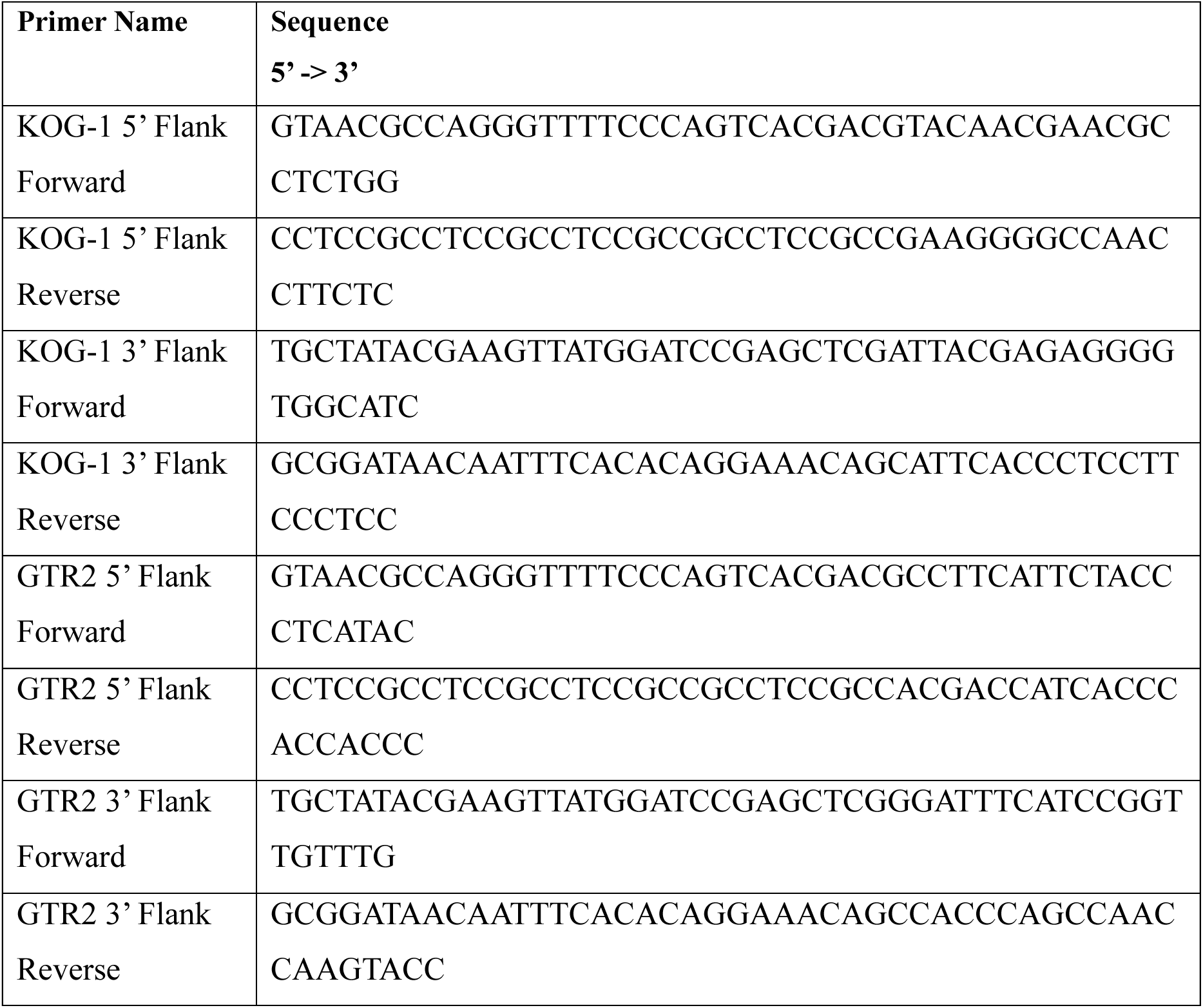
Primers.

### 2.2 Growth rate and period assays

Growth rates and circadian periods of the strains were assayed by growth on 30 cm glass race tubes as previously described (Adhvaryu et al., 2016). Race tubes contained maltose/arginine medium (MA) with 1X Vogel’s medium, 0.5% maltose, 0.01% arginine, 2% agar, and 100 µM choline to repair the choline defect in *chol-1* strains. Five replicate cultures of each strain were inoculated from conidiospores, germinated for 24 hours in constant light at 30°C, then transferred to constant darkness at 22°C and the growth fronts were marked daily on the underside of the tubes using red safelight. Growth rates and periods were calculated using in-house software as previously described (Lakin-Thomas, 1998). The data in Table S1 indicate that in some strains the *vta^ko^* mutation decreased growth rate by less than 10%, but the strains carrying KOG1 fusion proteins were unaffected. In all *vta^ko^* strains, the period was longer than wild type, as we have previously reported for the *vta^ko^* strain (Ratnayake et al., 2018). The KOG1-FLAG period was longer than wild type, but the KOG1-FLAG *vta^ko^*was significantly longer than the KOG1-FLAG strain without *vta^ko^*(Table S1).

### 2.3 TOR assays

The activity of TORC1 was assayed as previously described (Akhtari et al., 2025). Briefly, a FLAG epitope tag was knocked in at the endogenous S6 locus and the S6-FLAG was crossed into our standard strains. S6-FLAG strains were grown as described below and harvested by suction filtration, washed with water and flash frozen in liquid N_2_. Samples were ground to a powder in liquid N_2_, proteins were extracted in an SDS-containing buffer and immediately boiled and centrifuged. After protein assay, samples were loaded on SDS-PAGE gels containing phosphate-binding reagent PhosTag Acrylamide (Wako Chemicals) or Phos Binding Reagent Acrylamide (ApexBio) with MnCl_2_ to separate the phosphorylated forms of S6. Anti-FLAG antibody was used in immunoblotting to visualize three bands: S6, monophosphorylated S6, and di-phosphorylated S6 (S6-P). The ratio of the intensity of the S6-P band to the total of all three bands is reported as the S6-P ratio, and is used as a measure of TORC1 activity.

Cultures for TOR activity assay were maintained on agar slants containing 1x Vogel’s minimal medium, 2% glucose, 200 µM choline, and 2% agar. For high glucose tests (Fig. 1A), spores from 6-10 day old slant cultures were inoculated into 12-well plates with 2 ml of high glucose medium per well, containing 1xVogel’s, 2% glucose and 100 µM choline. Cultures were grown at 22°C in constant light for 48 hours. Each 48-h old mycelial mat was transferred to a small Petri dish with 5 ml high glucose medium for 4 hours to replenish the nutrients. Mats were then transferred to a fresh 5 ml plate of high glucose medium with the addition of amino acids for 1 h before harvest. For low glucose tests (Fig. 1B), spores were inoculated into 24-well plates with 1 ml of high glucose medium containing 1xVogel’s, 2% glucose,100 µM choline and 0.08% Tween 80. Cultures were grown at 22°C in constant light for 48 hours. The 48-hour old mycelial mats were transferred to a 150-ml flask (three mats per flask) with 50 ml of low-glucose medium containing 1X Vogel’s medium, 0.08% glucose and 100 µM choline and were shaken at 150 rpm for 24 h at 22°C in constant light. Glucose or amino acids were added to the flasks for 1 h before harvest.

**Fig. 1.**
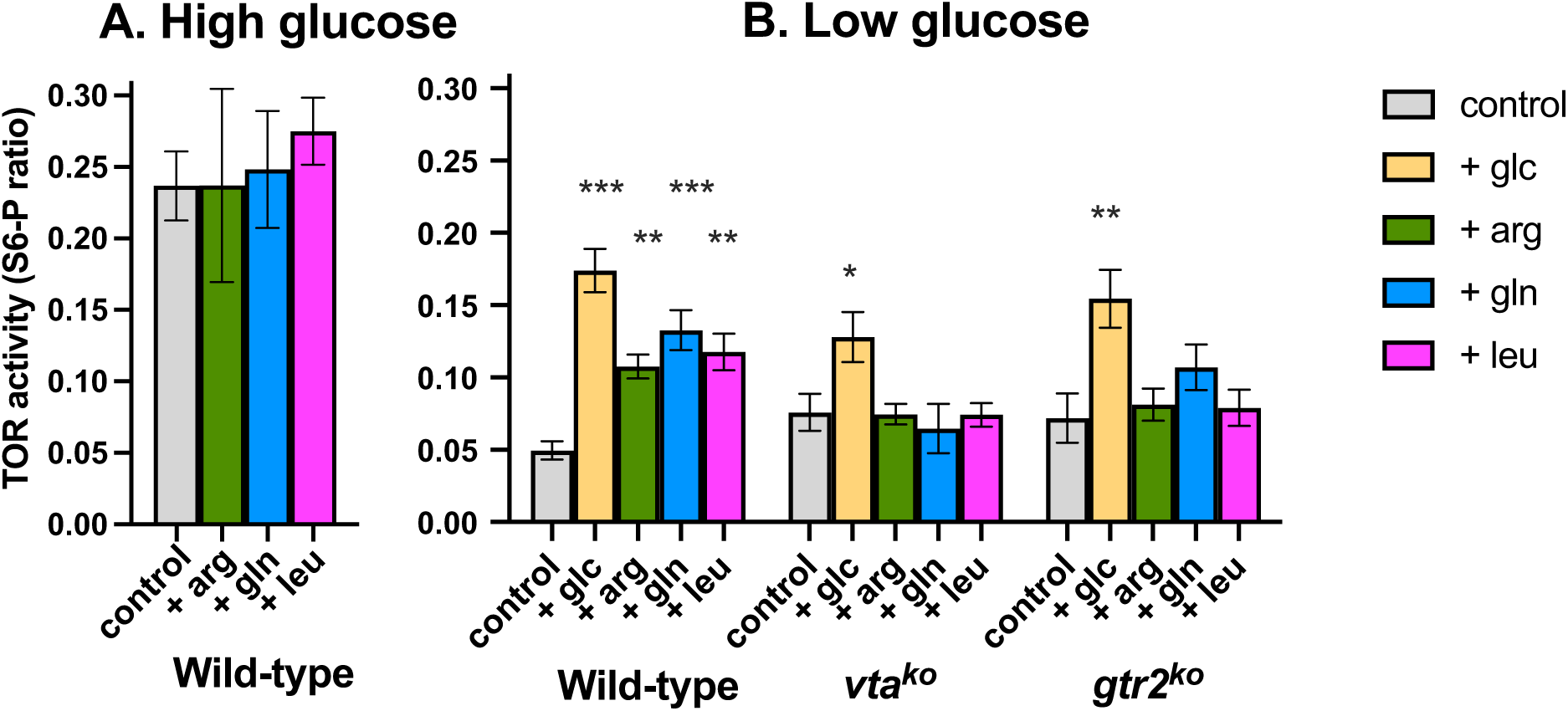
Activation of TOR by glucose and amino acids. TOR activity was assayed as the fraction of S6 protein in the fully phosphorylated state compared to total S6 protein. Data are plotted as mean ± SEM of 3-9 biological replicate experiments. Significance within each group was assessed by one-way ANOVA followed by Dunnett’s multiple comparison test. Significantly different from the control for that strain: * = p < 0.05; ** = p < 0.01; *** = p < 0.001. A. Cultures were grown in high glucose (2%) for 2 days and transferred to fresh high-glucose medium for 4 hours before addition of amino acids for one hour. B. Cultures were grown in high glucose for 2 days and then transferred to low glucose (0.08%) for 24 hours before addition of either glucose or amino acids for one hour.

Glucose was added as a water solution to low glucose cultures to a final 2% concentration. Amino acids were added as water solutions to final concentrations of 2.5 mM arginine, 5 mM glutamine, and 0.5 mM leucine. An equal volume of water was added to control cultures. Amino acid concentrations were chosen as the minimal concentration required to restore normal growth rates to the appropriate amino acid auxotrophic mutants, as reported in the literature.

### 2.4 Live Cell Microscopy

Spores were inoculated on 1.5% Phytagel (2% glucose, 1X Vogel’s salt and 100 µM choline) in Petri dishes and were grown at 22°C in constant light. When the mycelium had grown 3–5 cm from the point of inoculation, an agar block bearing the colony margin was cut from the edge of the colony and was inverted onto a 30 μL droplet of liquid medium on a 1.5 thickness 42 mm glass coverslip. This procedure puts the growing hyphae on the agar’s surface as close to the coverslip as possible (Hickey et al., 2004). Standard liquid glucose/arginine medium for observation contained 1X Vogel’s, 2% glucose, 5 mM arginine and 100 µM choline. Starvation medium omitted the glucose and arginine. Torin II inhibition medium was standard glucose/arginine with the addition of 500 nM Torin II. Cultures in starvation medium and Torin medium were incubated for at least 20 min before observation and imaging.

A Leica TCS SP8 Confocal microscope / Leica Dmi8 CS inverted stand was used, located at AOMF (Advanced Optical Microscopy Facility, University Health Network, Toronto, Canada). HC PL APO 63x/1.40 NA Oil immersion CS2 and HyD detectors were used for imaging. Line sequential setting was used and to take high resolution images the laser power was kept in the range of 1-4%. A pixel size of 2048×2048 was used. Averaging was established at 4 and the pinhole was 1 AU. Fiji Software was used to process images obtained.

### 2.5 Coimmunoprecipitation and mass spectrometry

Coimmunoprecipitation (Co-IP) was carried out as previously described (Eskandari et al., 2021) using Anti-FLAG M2 Affinity Gel beads (Sigma-Aldrich). FLAG-tagged strains were processed in parallel with untagged control strains. Frozen immunoprecipitates on beads were sent to SPARC BioCentre (SickKids Proteomics, Analytics, Robotics & Chemical Biology Centre, Hospital for Sick Children, Toronto, Canada) for mass spectrometry analysis. The results of mass spectrometry analysis were received as a Scaffold file (Scaffold 4.8., Proteome Software Inc). Candidates were found through the Uniprot database, and NCU numbers were identified and searched in FungiDB. Proteins with 95% probability in the total spectrum count that were only bound to the protein of interest and not control samples analyzed at the same time were chosen as binding partners, and candidates that were identified in two independent experiments bound to the same protein were listed as binding partners (Eskandari et al., 2021). Only binding partners with 10 or more total spectrum counts were included in Table S2.

## 3. Results

### 3.1 Activation of TOR by amino acids requires VTA and GTR2

To investigate the regulation of TOR by glucose and amino acids, we assayed TOR activity by measuring the phosphorylation of S6 ribosomal protein, which is a substrate of S6 kinase, which is in turn a substrate of TOR kinase. We FLAG-tagged the endogenous S6 gene and used anti-FLAG immunoblotting to assay the ratio of fully-phosphorylated S6 to total S6, as previously reported (Akhtari et al., 2025). The S6-FLAG was crossed into our standard laboratory strain, which will be referred to as the wild type. Strains carrying S6-FLAG in a *vta^ko^* or *gtr2^ko^* background were previously described (Akhtari et al., 2025). To remove rhythmic clock effects from the system, we grew cultures in constant light, which essentially “holds” the clock at one phase.

The amino acids arg, gln and leu were chosen from the literature on TOR activation in yeast and mammalian cells as prominent activators of TORC1 activity (González and Hall, 2017). Wild-type *N. crassa* does not require any amino acids for normal growth and in standard medium with adequate inorganic nitrogen and carbon sources it will not be deficient in amino acids. We found that when *N. crassa* cultures were grown and assayed in high glucose medium (2% glucose), the wild type did not respond to additional amino acid supplementation (Fig. 1A), indicating that glucose can fully activate TOR in the absence of additional amino acids. When wild-type cultures were held in low-glucose conditions (0.08% glucose for 24 hours), TOR activity was lower than in high glucose as expected and addition of glucose increased TOR activity (Fig. 1B). Addition of any of the three amino acids (without additional glucose) also increased TOR activity in the wild-type (Fig. 1B). In contrast, the *vta^ko^* and *gtr2^ko^* strains both responded to glucose addition but did not respond to amino acids (Fig. 1B). This indicates a requirement for the VTA gene product, as an EGO complex homolog, and GTR2 as a Rag-GTPase homolog, for amino acid regulation of TOR in *N. crassa*, but not for activation by glucose.

In low glucose medium (Fig. 1B), the levels of TOR activity in the controls without additional treatments appeared higher in the *vta^ko^* and *gtr2^ko^* mutants compared to wild type, but the differences are not statistically significant by one-way ANOVA (p > 0.19). Similarly, the levels of TOR activity after high glucose addition (Fig. 1B) are not significantly different among the three strains by one-way ANOVA (p > 0.20). We have previously reported (Akhtari et al., 2025) that TOR activity is rhythmic in strains growing in constant darkness on solid agar medium containing 0.5% maltose plus 0.01% arginine. The average TOR activity across several rhythmic cycles (the mesor) is also not significantly different among the wild type, *vta^ko^*and *gtr2^ko^* strains under these conditions (p > 0.10) (Akhtari et al., 2025). These results indicate that the VTA and GTR2 gene products do not appear to play a role in regulation of TOR activity in response to different carbon sources. We previously reported (Ratnayake et al., 2018) that the growth rate of the *vta^ko^* mutant, as measured by mass increase, does not fully respond to high levels of glucose as compared to the wild-type growth response. This indicates a lack of correlation between TOR activity (Fig. 1B) and growth rate (Ratnayake et al., 2018) in response to increased glucose availability.

### 3.2 Vacuolar localization of TORC1 and Rag-GTPase requires the EGO-like VTA protein

We have previously reported (Ratnayake et al., 2018) the vacuolar localization of the VTA gene product using a VTA-GFP fusion construct. Deletion of putative acylation sites abolishes this localization, suggesting that fatty acids anchor it to the membrane as these acyl groups anchor the yeast and mammalian homologs. To determine the relationships between VTA, GTR2 and the TOR complex, we constructed a KOG1-GFP fusion strain, as well as a GTR2-RFP fusion strain. The phenotypes of the tagged strains were assayed by growth on agar medium in race tubes, and the data in Table S1 and Fig. S1 indicate that the tagged proteins had only minor effects on growth rate and rhythmicity. The strains carrying *vta^ko^*and *gtr2^ko^* mutations displayed the characteristic damping of the conidiation rhythm previously reported (Eskandari et al., 2021; Li et al., 2011).

For microscopy, these strains were grown on solid Phytagel-containing medium and a block of the solid medium containing a sample of the growth front was inverted on a cover slip in a drop of liquid medium for live cell imaging. Based on our previous report of the vacuolar localization of VTA, and the vacuolar/lysosomal localization of TORC complexes in other organisms, we chose to concentrate our observations on older hyphae in which vacuoles could be seen. The VTA-GFP strain previously reported (Ratnayake et al., 2018) was imaged for comparison. As seen in Fig. 2 A, VTA was concentrated at the vacuolar membrane, outlining the margins of the vacuoles.

**Fig. 2.**
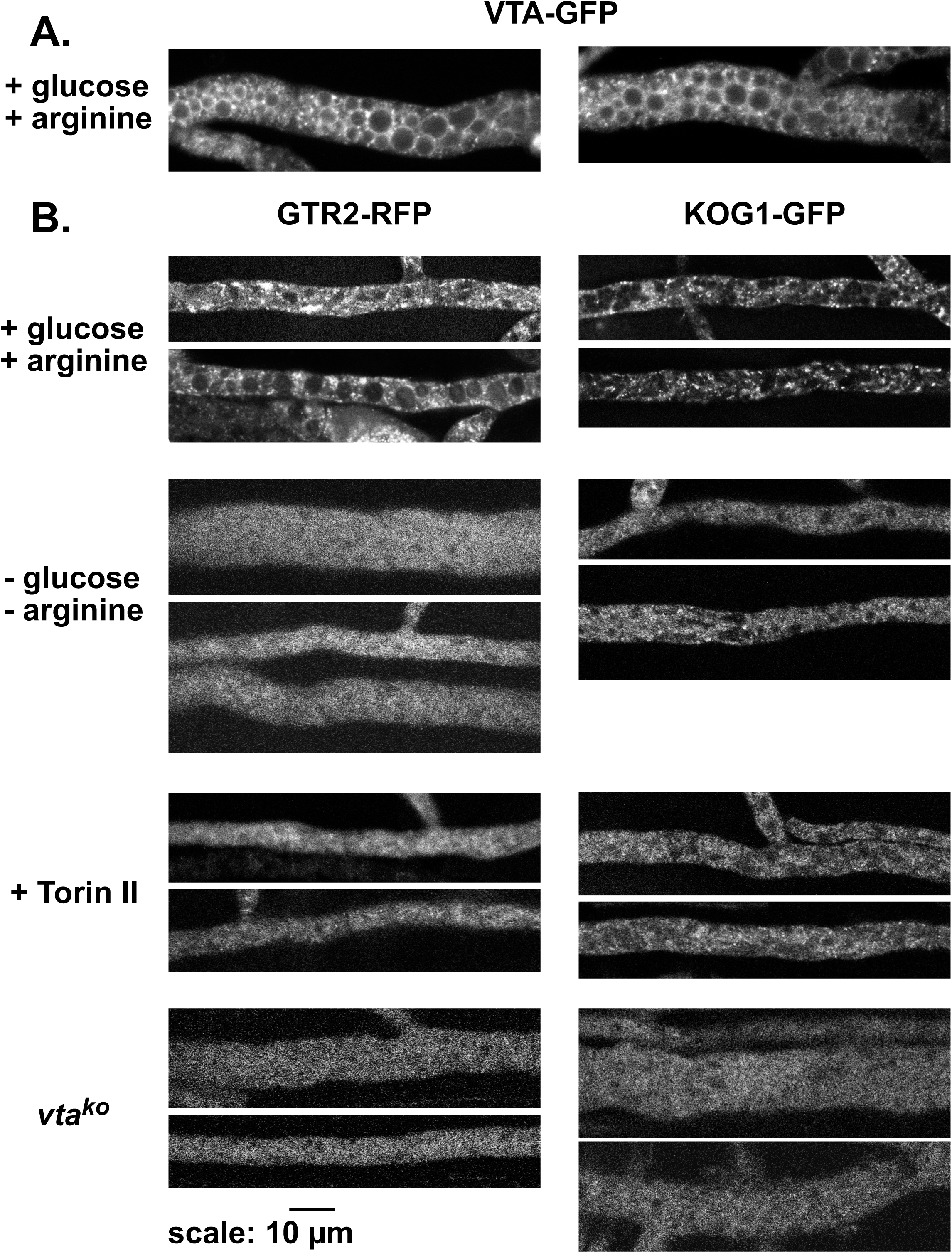
Localization of TOR complex proteins. GFP- and RFP-tagged proteins were imaged by live cell confocal microscopy. Two images from independent cultures are show for each condition. A. VTA-GFP in high glucose/arginine medium containing 2% glucose and 5 mM arginine (+ glucose/+ arginine). B. GTR2-RFP and KOG1-GFP strains imaged in high glucose/arginine (+ glucose/+ arginine), “starvation medium” with glucose and arginine omitted (- glucose/- arginine), high glucose/arginine with added 500 nM Torin II (+ Torin II), and strains carrying the *vta^ko^* mutation in high glucose/arginine (*vta^ko^*).

As seen in Fig. 2B, the GTR2-RFP and KOG1-GFP proteins were found associated with vacuoles in the rich (glucose/arginine) medium, but not as clearly associated with the membrane as VTA. These proteins appeared to cluster in punctate structures in the cytoplasm. After a short exposure (20 min) to starvation medium (-glucose, -arginine), the punctate structures were mostly lost and the proteins appeared dispersed in the cytoplasm. Torin II treatment for 20 min also dispersed the proteins, although some punctate structures could still be seen. In the *vta^ko^* mutant background, both proteins were evenly dispersed in the cytoplasm with no indication of structured localization.

These results indicate that the TOR complex (as localized by KOG1-GFP) is dispersed in the cytoplasm in the *vta^ko^* mutant. We have shown above (Fig. 1B) that baseline TOR activity levels are similar in the *vta^ko^* mutant as compared to the wild type on different carbon sources. Therefore, regulation of TOR activity by carbon source does not appear to require localization to the vacuole. In contrast, the loss of amino acid activation of TOR in the *vta^ko^* strain (Fig. 1B) and the cytoplasmic localization of KOG1-GFP in *vta^ko^* (Fig 2B) suggests that vacuolar localization of TOR is required for regulation by amino acids.

### 3.3 Composition of TOR complex depends on VTA protein

To identify the composition of the TORC1 complex in *Neurospora*, we constructed a FLAG-tagged KOG1 protein and crossed it into the wild-type background and the *vta^ko^*background. The GTR2-FLAG strains with wild type and *vta^ko^*background were previously described (Eskandari et al., 2021). FLAG-tagged proteins were precipitated from whole cell extracts prepared from strains carrying either the wild-type *vta* gene or *vta^ko^* and binding partners were determined by mass spectrometry. Peptide counts and spectrum counts for top binding partners are presented in Table S2, and binding partners are displayed graphically in Fig. 3.

**Fig. 3.**
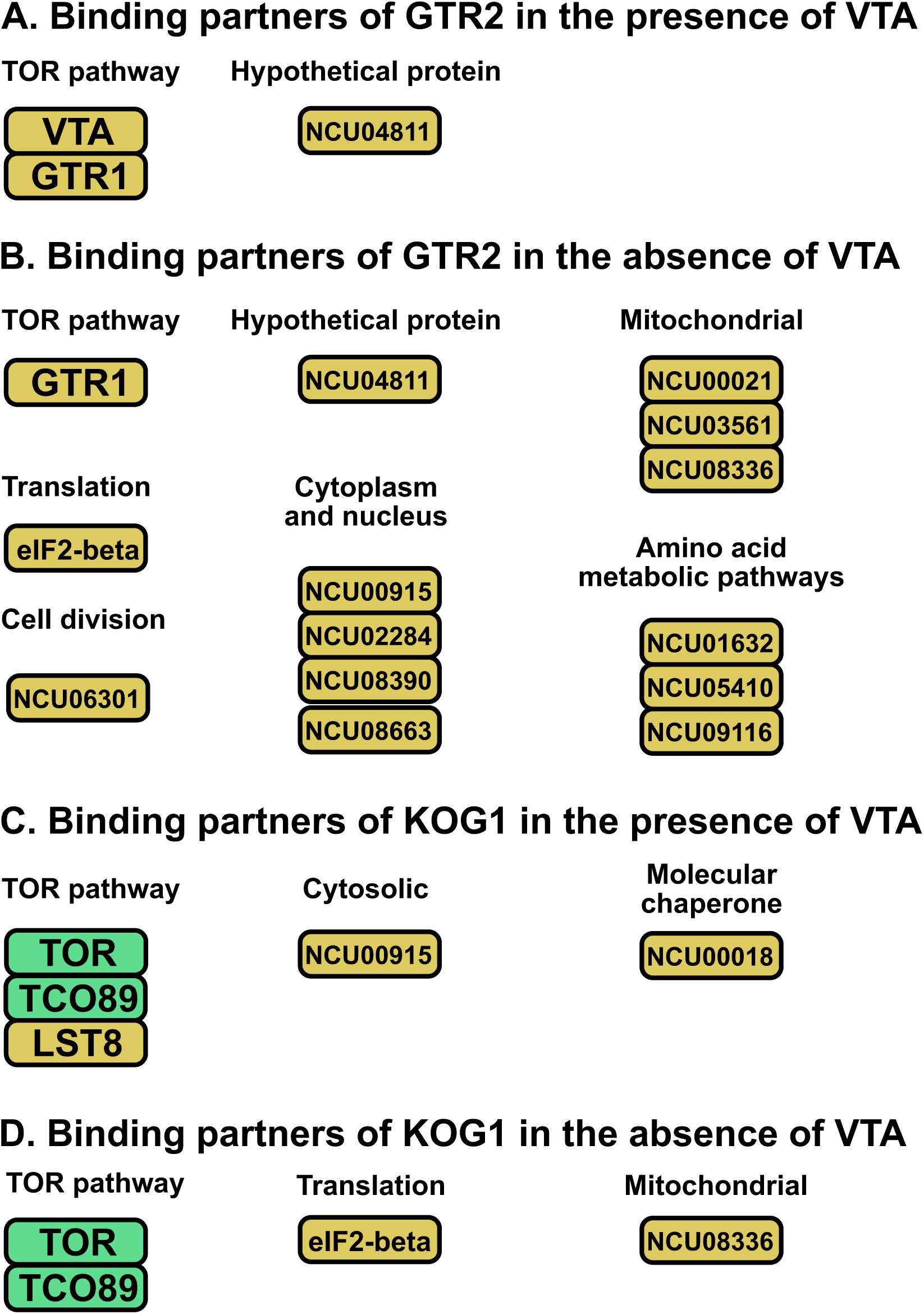
Binding partners of GTR2 and KOG1. FLAG-tagged GTR2 and KOG1 were immunoprecipitated from strains carrying either wild-type *vta* or the *vta^ko^* null allele and the binding partners were identified by mass spectrometry. Partners with total peptide counts above 90 from two independent experiments are in green; partners with counts between 10 and 90 are in yellow. A. GTR2-FLAG in wild-type *vta* background. Data are from (Eskandari 2021). B. GTR2-FLAG in *vta^ko^* background. C. KOG1-FLAG in wild-type *vta* background. D. KOG1-FLAG in *vta^ko^* background.

We previously reported (Eskandari et al., 2021) that GTR2 binds, as expected, to GTR1 (NCU01099), the homolog of the other Ras-GTPase found in the EGO complex in yeast, and VTA, thereby anchoring the Ras-GTPases to the vacuolar membrane (Fig 3A). In the *vta^ko^* background, GTR2 is still bound to GTR1, but is also found in association with a number of cytoplasmic proteins (Fig. 3B), suggesting non-specific binding due to cytoplasmic localization. This is consistent with the uniform cytoplasmic localization seen by microscopy in the *vta^ko^* background (Fig. 2B). In both *vta^+^* and *vta^ko^* backgrounds, an unidentified hypothetical protein NCU04811 is found in the complex.

KOG1 protein was found in a complex with the expected components TOR kinase (NCU05608) and LST8 (NCU04281) (Fig. 3C). A fourth TOR complex component TCO89 (NCU04795) found in fungi (Reinke et al., 2004) was also associated with KOG1. In the absence of VTA, TOR, TCO89 and KOG1 remained in a complex but LST8 was lost (Fig. 3D).

## 4. Discussion

We have observed increased activity of TOR in response to added glucose in the *vta^ko^* mutant (Fig. 1B) but a failure of growth rate (in liquid medium) to fully respond to added glucose in this mutant (Ratnayake et al., 2018). We previously reported (Akhtari et al., 2025) that growth on solid agar medium, measured as either linear growth rate or mass increase, also does not correlate with TOR activity. This may indicate that TOR activity does not always directly drive growth rate, suggesting a complex feedback system between downstream effects of TOR and upstream modulators of TOR activity.

Our localization results (Fig. 2) may indicate some difference between *N. crassa* and localizations reported for yeast. As we reported previously (Ratnayake et al., 2018), VTA is found evenly distributed around the vacuolar membrane (Fig. 2A), consistent with its role as an integral membrane protein associated with the outer lipid leaflet through covalently attached acyl groups (Ratnayake et al., 2018). In conditions of high nutrition (glucose plus arginine), both GTR2 and KOG1 are found in punctate structures near the vacuoles (Fig 2B). This structure is partially disrupted in starvation conditions (minus glucose and arginine) and also during inhibition by Torin II (Fig. 2B). This suggests that active TOR is required for the association of the TORC1 complex and the amino acid sensing Ras-GTPase complex with vacuolar components. In the *vta^ko^* null background, both GTR2 and KOG1 appear uniformly cytoplasmic, identifying the VTA-containing EGO-like vacuolar complex as the anchor for vacuolar localization.

In yeast, Kog1 moves from its localization at the vacuolar membrane to a single body near the edge of the vacuole in the glucose starvation condition in a rapid and reversible process (Hughes Hallett et al., 2015). During starvation AMPK activates Snf1, which phosphorylates glutamine-rich prion-like motifs in Kog1. In a high number of organisms lacking TSC1/2, such as yeast and *C. elegans*, the Snf1-dependent phosphorylation sites were present. Therefore, it appears that regulation of TORC1 in early eukaryotes is by Kog1-body formation, while higher eukaryotes use TSC1/2. In *N. crassa* TSC1/2 is not expressed (Diernfellner et al., 2019). Our results (Fig. 2B) show punctate structures that may be similar to Kog1 bodies in the presence of high nutrition, but these are disrupted by starvation or Torin II inhibition, suggesting a somewhat different regulation of TORC1 activity in *N. crassa* compared to yeast.

Our co-IP results (Fig. 3) support the localizations seen in Fig. 2. A comparison of the GTR2-FLAG *vta^ko^* mass spectrometry results (Fig. 3B) with previous results (Eskandari et al., 2021) indicates that GTR2 is still binding to the two major proteins that it bound in the presence of VTA, including GTR1 (NCU01099) and a hypothetical protein NCU04811. However, we found many more binding partners for GTR2 in the absence of VTA. We hypothesize that the dissociation of GTR2 from VTA does not affect the interactions of GTR2, GTR1, and the hypothetical protein. However, it does dissociate the GTR2-GTR1 complex from the vacuolar membrane, which increases the interaction of GTR2 with numerous other cytoplasmic proteins. This is also consistent with our microscopy observations, as we see that in the absence of VTA there is no localization of the GTR2 protein around the vacuole and the expression is cytoplasmic.

The most abundant common binding partner shared by KOG1-FLAG and KOG-1-FLAG *vta*^ko^ is NCU05608, identified as Serine/Threonine protein kinase TOR. This is expected since KOG1 is known as a key regulator of TORC1 activation and directly binds to the TOR protein. It was reported that amino acid starvation does not affect the localization or the direct interaction of mTOR with Raptor in mammalian cells (Yadav et al., 2013). This is consistent with our mass spectrometry results, as we can see that KOG1, a homolog of Raptor, is still bound to TOR protein in the presence and absence of VTA. The next most abundant protein is NCU04795, identified as TCO89. TCO89 is found as a component of the TORC1 complex in fungi (Nicastro et al., 2025; Reinke et al., 2004) and functions as a tether connecting TORC1 to the RAG GTPases (Nicastro et al., 2025). We find TCO89 is still in association with TOR and KOG1 in the absence of VTA, but we do not find the Rag GTPases GTR1 and GTR2 bound to TORC1 in the presence or absence of VTA. This may indicate that in *N. crassa* the vacuolar localization of the Rag GTPases in the presence of VTA is required for localization of TORC1 on the vacuole, and GTR1/GTR2 do not associate with TORC1 without this localization, unlike the findings in yeast (Nicastro et al., 2025).

In this paper, our goal was to add to our understanding of the TOR-related functions of the VTA and GTR2 gene products in *N. crassa*, to further explore their roles in regulating circadian rhythmicity. The current model for the *N. crassa* circadian oscillator, called the FRQ/WCC TTFL (transcription/translation feedback loop), comprises interlocked negative and positive feedback loops that include the clock genes *frq, wc-1* and *wc-2* (Baker et al., 2012; Bell-Pedersen, 2000; Lakin-Thomas et al., 2011). WC-1 and WC-2 proteins heterodimerize to form the White-Collar Complex (WCC). When WCC is active the levels of *frq* mRNA will increase and FRQ protein is translated. The FRQ protein then will negatively regulate its own transcription by inhibiting the activity of WCC and lead to a decrease in *frq* mRNA levels. The resulting rhythmic activity of the WC complex drives the rhythmic outputs of the clock system such as conidiation.

Although most investigation on the *N. crassa* circadian system has been dedicated to the FRQ/WCC feedback loop, it has been known for many years that rhythmic conidiation can be seen in the absence of a functional *frq* gene. Numerous examples indicating the presence of FRQ-less rhythms in various growth conditions and genetic backgrounds have been reported (Lakin-Thomas et al., 2011; Ratnayake et al., 2018). There must be an oscillator that regulates rhythmicity in the absence of FRQ/WCC function but the identity of this oscillator is unknown. Our lab is studying this FRQ-less oscillator (FLO) and its association with the FRQ/WCC TTFL. To identify components of the FLO, we used a mutagenesis screen in an FRQ-less strain and identified a mutation we named uv90 that abolishes FRQ-less rhythmicity of conidiation (Li et al., 2011). When we studied the phenotype of this uv90 mutation in a FRQ wild-type strain, we found that it dampened the conidiation rhythm as well as the amplitude of the FRQ protein. In assaying responses of the clock to environmental signals (light pulses and heat pulses) we found that the responses of the uv90 mutant were altered, indicating a decrease in the amplitude of the oscillator’s cycle. The uv90 mutation, therefore, identifies a gene necessary for robust FRQ-less rhythmicity and for maintaining the amplitude of the entire circadian system and the normal functioning of the FRQ/WCC TTFL when FRQ is present.

We mapped the uv90 mutation and identified the gene as a member of the LAMTOR family, similar to p18/LAMTOR1 in mammals and EGO1/Meh1p/Gse2p in yeast, which are components of the Ragulator and EGO complexes that anchor TORC1 to the lysosomal/vacuolar membrane (Ratnayake et al., 2018). We therefore named the gene product VTA, for vacuolar TOR-associated protein. Coimmunoprecipitation of a VTA-FLAG strain identified GTR2 as a top binding partner (Ratnayake et al., 2018). Reciprocal coimmunoprecipitation of GTR2-FLAG indicated that VTA and components of the TOR pathway also coimmunoprecipitated with GTR2 (Eskandari et al., 2021). The phenotype of a *gtr2^ko^*null mutant is very similar to the *vta^ko^* null mutant, dampening the conidiation rhythm in the FRQ wild-type background, dampening the FRQ protein rhythm, and abolishing FRQ-less conidiation rhythmicity (Eskandari et al., 2021).

We have assayed the effects of *vta^ko^* and *gtr2^ko^*on the rhythm of TORC1 activity and found that VTA and GTR2 are not required to maintain baseline TOR activity but are required for maintaining TOR rhythmicity (Akhtari et al., 2025). In the *vta^ko^*and *gtr2^ko^* knockout strains, the TOR rhythm gradually damps out but TOR activity remains at a mean level similar to the wild type. This suggests that rhythmicity of TOR activity is required to maintain circadian rhythmicity, and not simply that active TOR is a requirement. The question becomes, what functionality of the TOR pathway is lost in these mutants? Our results in Fig 1 indicate that it is the ability to respond to amino acids that is missing, not the response to increased carbon source (glucose). This focusses attention on the amino acid sensing complex as a potential regulator of rhythmicity. We have previously proposed (Lakin-Thomas, 2023) that feedback loops inherent in the TOR pathway could form the basis for an oscillator driven by negative feedback of TOR on its own activity. The depletion of amino acids by protein synthesis, and the supply of amino acid from autophagy which is inhibited by TOR, could provide the negative feedback for such an oscillator. Our current results strengthen this as a possibility.

## Declaration of Competing Interests

The authors declare that they have no known competing financial interests or personal relationships that could have appeared to influence the work reported in this paper.

## Acknowledgements

This work was supported by Natural Sciences and Engineering Research Council of Canada Discovery Grant number RGPIN-2017-05664.

Technical assistance by Shahrzad Rahmani, Arghavan Sammak and Ethan Sooklal is gratefully acknowledged. We thank the Fungal Genetics Stock Center, Kansas State University, for plasmids and strains.

## Supplementary material

Supplementary Figure S1 and Tables S1 and S2 can be found in the accompanying file.

## Notes

### Competing Interest Statement

The authors have declared no competing interest.

