## Supplemental Information for "EGO-like complex regulates TOR (Target of Rapamycin) activity and localization in *Neurospora*"

### Supplementary Information

**Fig. S1. Growth phenotypes of strains**
Two representative race tubes from each set of 5 replicate cultures are shown. All strains carry the *ras*$^{bd}$, *csp-1* and *chol-1* mutations in addition to the indicated mutations. Cultures were photographed from above for easier visualization of the banding pattern so daily growth marks on the underside are not visible; the white bar on the wild type image indicates the approximate extent of growth in 24 hours. Green colouration is from food colouring added to the medium for improved visibility.

**Table S1. Growth rates and periods of strains**

**Table S2. Co-immunoprecipitation and mass spectrometry**

**Fig. S1**

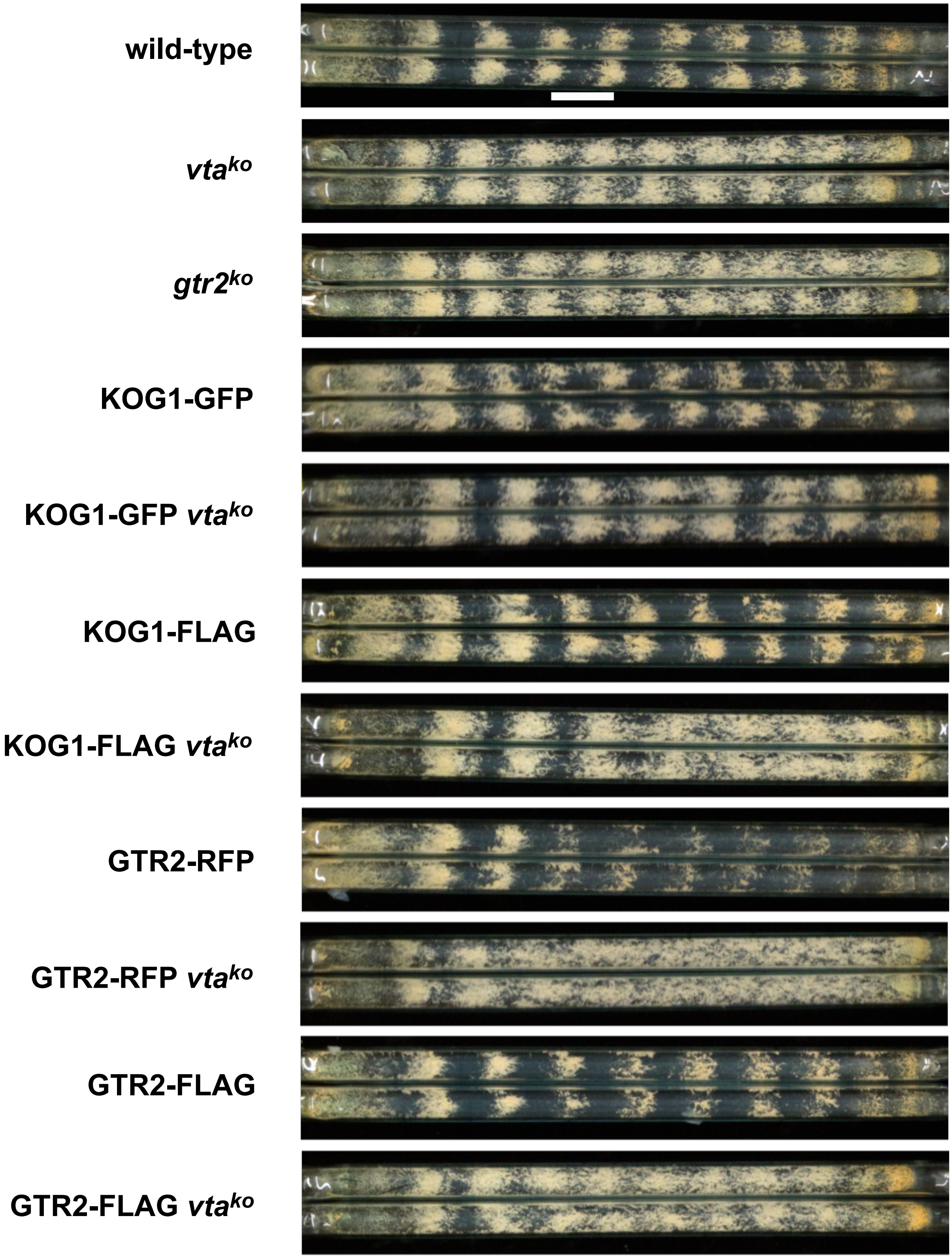
wild-type
*vta*$^{ko}$
*gtr2*$^{ko}$
KOG1-GFP
KOG1-GFP *vta*$^{ko}$
KOG1-FLAG
KOG1-FLAG *vta*$^{ko}$
GTR2-RFP
GTR2-RFP *vta*$^{ko}$
GTR2-FLAG
GTR2-FLAG *vta*$^{ko}$

**Table S1. Growth rates and periods of strains**

| **Strain** | **Growth rate (mm/h)** | **Period (h)** |
|---|---|---|
| wild-type | 1.14 ± 0.01 | 20.12 ± 0.31 |
| *vta*$^{ko}$ | 1.09 ± 0.01* | 22.12 ± 0.12*** |
| *gtr2*$^{ko}$ | 1.04 ± 0.02** | 22.07 ± 0.27** |
| KOG1-GFP | 1.07 ± 0.01** | 20.84 ± 0.28 |
| KOG1-GFP *vta*$^{ko}$ | 1.15 ± 0.01 | 21.30 ± 0.10** |
| KOG1-FLAG | 1.14 ± 0.01 | 21.55 ± 0.04** |
| KOG1-FLAG *vta*$^{ko}$ | 1.13 ± 0.01 | 24.09 ± 0.43*** |
| GTR2-RFP | 1.13 ± 0.003 | 20.03 ± 0.17 |
| GTR2-RFP *vta*$^{ko}$ | 1.07 ± 0.01*** | 26.03 ± 0.56*** |
| GTR2-FLAG | 1.20 ± 0.03 | 20.21 ± 0.16 |
| GTR2-FLAG *vta*$^{ko}$ | 1.08 ± 0.01** | 22.68 ± 0.75* |

All strains carry the *ras*$^{bd}$, *csp-1* and *chol-1* mutations in addition to the indicated genotypes. Strains were grown in race tubes on media containing 100µM choline to repair the *chol-1* deficiency. Data are reported as mean ± S.E.M for N = 5 race tubes. Means were compared to wild-type by Student's unpaired t-test. *: $p < 0.05$; **: $p < 0.01$; ***: $p < 0.001$

### Table S2

#### Co-immunoprecipitation and mass spectrometry

| Locus ID | Gene product | MW (kDa) | Exp. 1 | | Exp. 2 | | Total counts |
|---|---|---|---|---|---|---|---|
| | | | Peptides | Counts | Peptides | Counts | |
| **Binding Partners of KOG1-FLAG** | | | | | | | |
| NCU05608 | Serine/threonine-protein kinase TOR | 278 | 2 | 187 | 2 | 11 | 198 |
| NCU04795 | TCO89 | 76 | 2 | 91 | 2 | 12 | 103 |
| NCU00915 | aspartyl-tRNA synthetase | 65 | 2 | 4 | 2 | 29 | 33 |
| NCU04281 | LST8 | 36 | 2 | 17 | 2 | 3 | 20 |
| NCU00018 | CDC48 | | 2 | 7 | 2 | 11 | 18 |
| **Binding Partners of KOG1-FLAG *vta* $^{ko}$** | | | | | | | |
| NCU05608 | Serine/threonine-protein kinase TOR | 278 | 2 | 143 | 2 | 7 | 150 |
| NCU04795 | TCO89 | 76 | 2 | 82 | 2 | 12 | 94 |
| NCU04640 | eIF-2 beta | 32 | 2 | 5 | 2 | 17 | 22 |
| NCU08336 | Succinate dehydrogenase | 71 | 2 | 5 | 2 | 8 | 13 |
| **Binding Partners of GTR2-FLAG *vta* $^{ko}$** | | | | | | | |
| NCU09116 | Aromatic aminotransferase Aro8 | 62 | 2 | 26 | 2 | 36 | 62 |
| NCU05410 | Acetylornithine transaminase | 33 | 2 | 14 | 2 | 24 | 38 |
| NCU01632 | Pentafunctional arom polypeptide | 170 | 2 | 4 | 2 | 30 | 34 |
| NCU04640 | eIF-2 beta | 32 | 2 | 11 | 2 | 20 | 31 |
| NCU01099 | Gtr1 RagA | 45 | 2 | 21 | 2 | 5 | 26 |
| NCU08390 | N-acetyltransferase domain-containing protein | 27 | 2 | 6 | 2 | 11 | 17 |
| NCU03561 | Mitochondrial carrier | 33 | 2 | 10 | 2 | 5 | 15 |
| NCU04811 | Uncharacterized protein | 32 | 2 | 11 | 2 | 3 | 14 |
| NCU02284 | nucleolar ATPase Kre33 | 118 | 2 | 3 | 2 | 11 | 14 |
| NCU00915 | Aspartyl-tRNA synthetase | 65 | 2 | 6 | 2 | 7 | 13 |

| | | | | | | | |
|---|---|---|---|---|---|---|---|
| NCU08663 | nonsense-mediated mRNA decay protein 3 | 58 | 2 | 4 | 2 | 8 | 12 |
| NCU08336 | Succinate dehydrogenase flavoprotein subunit | 71 | 2 | 6 | 2 | 6 | 12 |
| NCU06301 | CENP-V/GFA domain-containing protein | 18 | 2 | 6 | 2 | 5 | 11 |
| NCU00021 | Clustered mitochondria protein homolog | 142 | 2 | 3 | 2 | 8 | 11 |

Binding partners of each FLAG-tagged bait protein are listed in descending order of total spectrum counts. Proteins with fewer than 10 total counts are not included. An unrelated FLAG-tagged strain and a strain without any FLAG tag were used as controls in each experiment. NCU number represents the ID of the gene in the *N. crassa* genome database (FungiDB). Results of two independent experiments are reported.
